# Electrical synapses and transient signals in feedforward canonical circuits

**DOI:** 10.1101/394593

**Authors:** Tuan Pham, Julie S. Haas

## Abstract

As information about the world traverses the brain, the signals exchanged between neurons are passed and modulated by synapses, or specialized contacts between neurons. While neurotransmitter-based synapses tend to be either relay excitatory or inhibitory pulses of influence on the postsynaptic neuron, electrical synapses, composed of plaques of gap junction channels, are always-on transmitters that can either excite or inhibit a coupled neighbor. A growing body of evidence indicates that electrical synapses, similar to their chemical counterparts, are modified in strength during physiological neuronal activity. The synchronizing role of electrical synapses in neuronal oscillations has been well established, but their impact on transient signal processing in the brain is much less understood. Here we constructed computational models based on the canonical feedforward neuronal circuit, and included electrical synapses between inhibitory interneurons. We provided discrete closely-timed inputs to the circuits, and characterize the influence of electrical synapses on both the subthreshold summation and spike trains in the output neuron. Our simulations highlight the diverse and powerful roles that electrical synapses play even in simple circuits. Because these canonical circuits are represented widely throughout the brain, we expect that these are general principles for the influence of electrical synapses on transient signal processing across the brain.

**Author Summary:** The role that electrical synapses play in neural oscillations, network synchronization and rhythmicity is well established, but their role neuronal processing of transient inputs is much less understood. Here we used computational models of canonical feedforward circuits and networks to investigate how the strength of electrical synapses regulates the flow of transient signals passing through those circuits. We show that because the influence of electrical synapses on coupled neighbors can be either inhibitory or excitatory, their role in network information processing is heterogeneous.. Because of the widespread existence of electrical synapses between interneurons as well as a growing body of evidence for their plasticity, we expect such effects play a significant role in how the brain processes transient inputs.

## Introduction

Electrical synapses are prevalent across many brain regions, including thalamus, hypothalamus, cerebellum, and the neocortex [1, 2]. In contrast to neurotransmitter-based synapses, electrical synapses are an always-on mode of intracellular communication. Because signals cross two cell membranes, the net effect of an electrical synapse is that of a lowpass filter: spikes are heavily attenuated, while longer events, such as bursts, UP states, and the depolarizations that lead to spikes, are more readily shared between cells. Further, electrical synapses can exert either inhibitory or excitatory effects on a coupled neighbor, by increasing leak at rest or by transmitting activity such as depolarizations or spikelets. A growing body of work has demonstrated ways in which electrical synapses can be modulated or modified by either synaptic [3-8] or spiking [9, 10] forms of neuronal activity.

The role of electrical synapses in neuronal signal processing has mainly been explored in terms of their contributions to or regulation of synchrony of ongoing oscillations [11-17]. Studies focusing on the influence of electrical synapses on transient signals as they traverse the brain are fewer, but hint at specific and potentially powerful roles. For instance, propagation of spike afterhyperpolarizations through electrical synapses acts to reset and desynchronize regular firing in coupled cerebellar Golgi neurons [18]. Electrical synapses accelerate timing of spikes elicited near threshold in coupled thalamic reticular neighbors by tens of ms [19, 20], and similarly in coupled cerebellar basked cells, electrical synapses enhance and accelerate recruitment for coincident or sequential inputs [21]. Axonal gap junctions between neurons in the fly visual stream aid efficient encoding of the axis of rotation [22]. Our previous work focused on the impact of electrical synapses on transient signals in the thalamacortical relay circuit, showing that electrical coupling between inhibitory neurons lead to increased separation of disparately-timed input while facilitating fusion of closely-timed inputs [23].

In order to generalize a role for electrical synapses and variations in their strength in neuronal information processing, here we considered the canonical microcircuit, wherein two principal neurons, connected by an excitatory synapse, are also connected by disynaptic feedforward inhibition (Fig.1A_1_)[24]. This circuit motif reappears through the brain in areas ranging from the hippocampal CA1 pyramidal neurons [25], somatosensory L4 cortical neurons receiving inputs from the ventrobasal complex [26], and the cortical translaminar inhibitory circuit [27] (Fig. 1A_2-4_) Starting with a canonical circuit, we progressively expanded models to include interneurons coupled by electrical synapses in both circuit and network configurations. We provided these models with closely timed inputs, in order to determine how the embedded electrical synapse influences the summation or integration of those inputs at the output stage of the model. Our simulations demonstrate that electrical synapses between interneurons of the canonical circuit enable a high degree of specificity and diversity of processing of transient signals for both subthreshold activity and network activity. Because electrical synapses are widespread throughout the mammalian brain, we expect that these are principles that apply widely to neuronal processing of newly incoming information as it passes through the brain.

**Figure 1.**
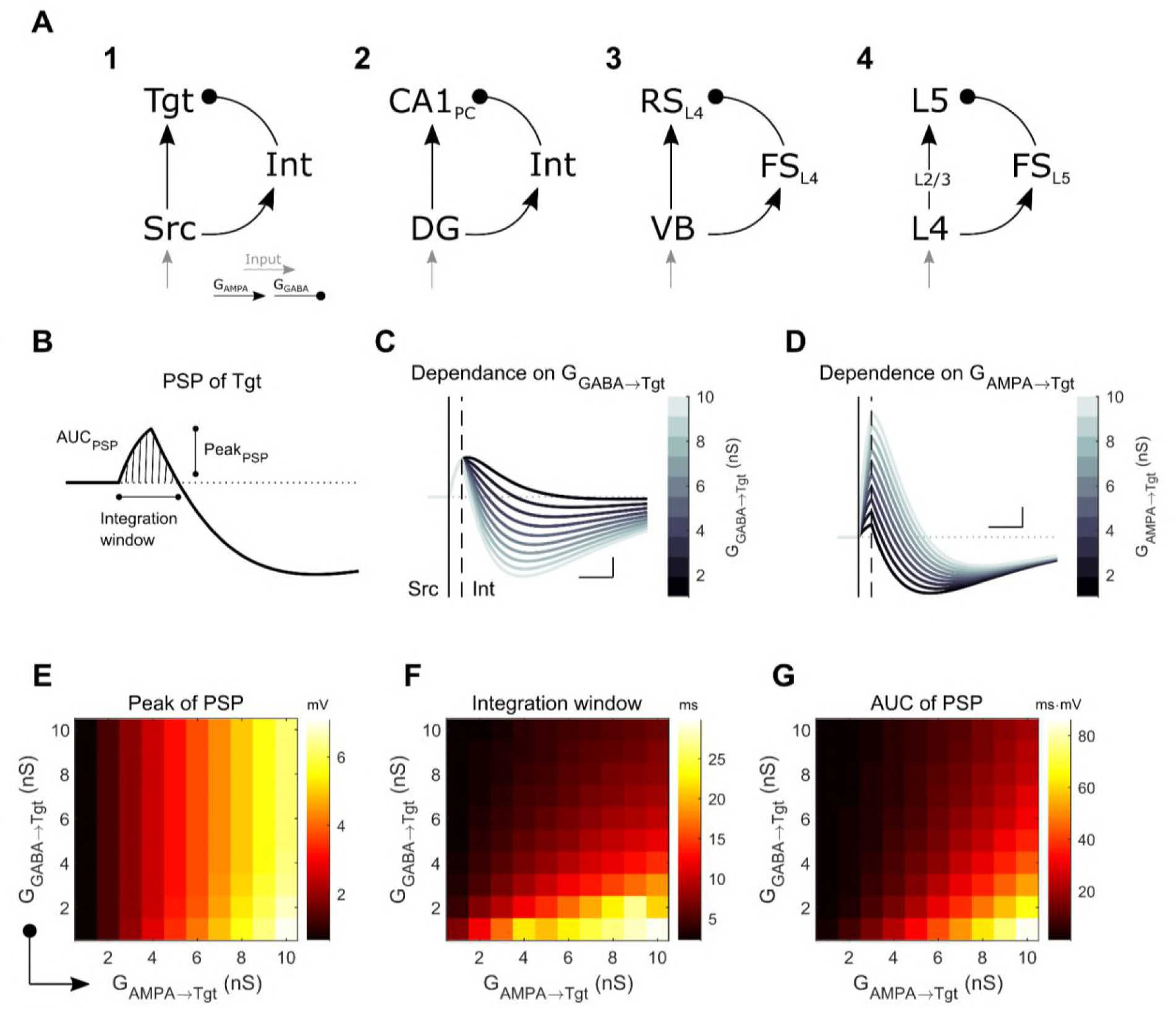
Model 1a: the canonical model of feedforward inhibition. **A**: Three-cell circuit model (A1) with feedforward disynaptic inhibition between excitatory source (Src) and target (Tgt) neurons. This canonical model represents those found in for example (A2) the hippocampal circuit, between dentate gyrus (DG) and CA1 cells [25]; (A3) from thalamic VB relay neurons to regular spiking cells in the somatosensory thalamocortical circuit [26]; and (A4) the cortical translaminar inhibitory circuit [27]. **B**: Example compound subthreshold postsynaptic membrane potential (PSP) in the Tgt neuron following a spike in Src, and the quantifications (PSP peak, integration window, and area under the PSP curve (AUC)) used throughout the text. **C**: Effect of different inhibitory strengths G_GABA→Tgt_ on the compound PSP in Tgt; G_AMPA→Tgt_ was 3nS. **D**: Effect of varied G_AMPA→Tgt_ on the compound PSP of Tgt; G_GABA→Tgt_ was 6nS. For both C and D, scale bar is 1mV, 5ms; the vertical straight line and dashed line mark the spike times of Src and Int, respectively. **E-F**: Combined effects of both excitatory and inhibitory synaptic strengths towards the peak, duration of the integration window and AUC of the positive portion of the compound PSP in Tgt.

## Results

### 1. Subthreshold integration in a canonical circuit with electrical synapses

We started our inquiry by creating a three-cell circuit that represents the canonical microcircuit: two excitatory neurons, with an interneuron providing feedforward inhibition (Fig. 1A). This model (Model 1a) produces a compound postsynaptic potential (PSP) in the target (Tgt) neuron that is a sum of a purely excitatory PSP from the source (Src) neuron and an inhibitory PSP arriving with a delay from the inhibitory interneuron (Int) (Fig. 1B). The features of the PSP – its height, its total excitation, and the duration of its excitation window, all of which in combination may determine whether Tgt will spike given sufficient input –depend predictably on the strength of the inhibitory PSP arriving from Int (Fig. 1C). Increases in G_GABA→Tgt_ limit the integration window (Fig. 1F) and the area under excitatory portion of the PSP (Fig. 1G). Thus, the interneuron limits the overall excitation and possible triggering of an action potential in Tgt.

To understand how Tgt might sum input from multiple sources, our next step towards building larger models was to include two Src neurons, which both synapsed onto a common Int and a common Tgt (Model 1b, Fig. 2A). We provided both Src neurons with brief inputs sufficient to evoke single spikes in the Src neurons, while varying the time delay between the inputs Δt_inp_. From these simulations, we observed that the inhibition from the common Int limited summation of the two Src signals in Tgt (Fig. 2B) in two ways. Input timing differences of Δt_inp_ >1 ms led to sharp decreases in PSP peaks in Tgt (Fig.2C), as the inhibition from Int initiated by one Src decreased summed response with the second Src in Tgt. Summation of the Src signals in Int also accelerated the Int spike (Fig. 2B), thereby limiting the integration window in Tgt (Fig. 2D). For delays between two inputs Δt_inp_ longer than ∼4 ms, a second input was consistently prevented from summation with the first input by the feedforward inhibition arriving as a result of the first input (Fig. 2C-E). Additionally, we noted that decreased excitability of the interneuron, in this case by providing less holding current to Int but which could arise from increased leakage in electrically coupled networks, shifted PSP summation, integration window and AUC incrementally towards larger values (Fig. 2C-E). These results provide a baseline of expectations for a circuit with globally strong feedforward inhibition, for example one that includes interneurons that are strongly coupled.

**Figure 2.**
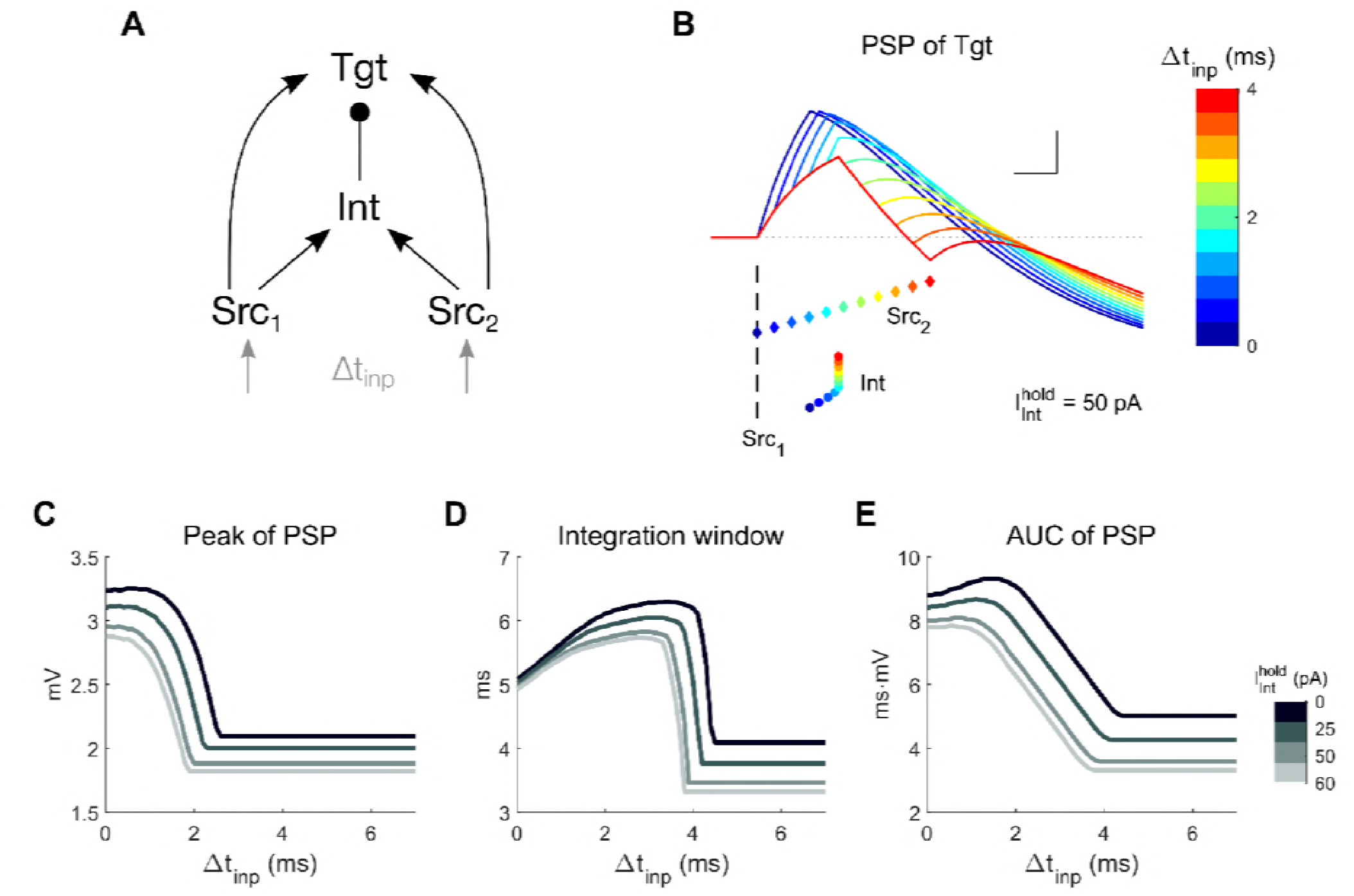
Model 1b: Two Src neurons synapse onto a common Int and Tgt. **A**: Model schematic; each Src neuron receives its own input, with timing difference between the two inputs Δt_inp_. **B**: Example PSP in Tgt when the inputs arrive within 4ms of each other, with a color code representing different values of Δt_inp_. Spike times are shown below for Src_1_ (vertical dashed line ---), Src_2_ (colored diamonds ♦) and Int (colored circles •). Scale bar is 1mV, 1ms. **C-E**: Effect of varied interneuron excitability 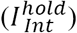 on Tgt PSP properties.

Next, we expanded the circuit model to include two inhibitory Int neurons coupled by an electrical synapse (Model 2a, Fig. 3A). We limited the range of strength of the electrical synapse to vary between 0 (uncoupled) and a coupling coefficient of ∼0.3, which represents common strengths found in the thalamus [10, 28] and cortex [29-31].

**Figure 3.**
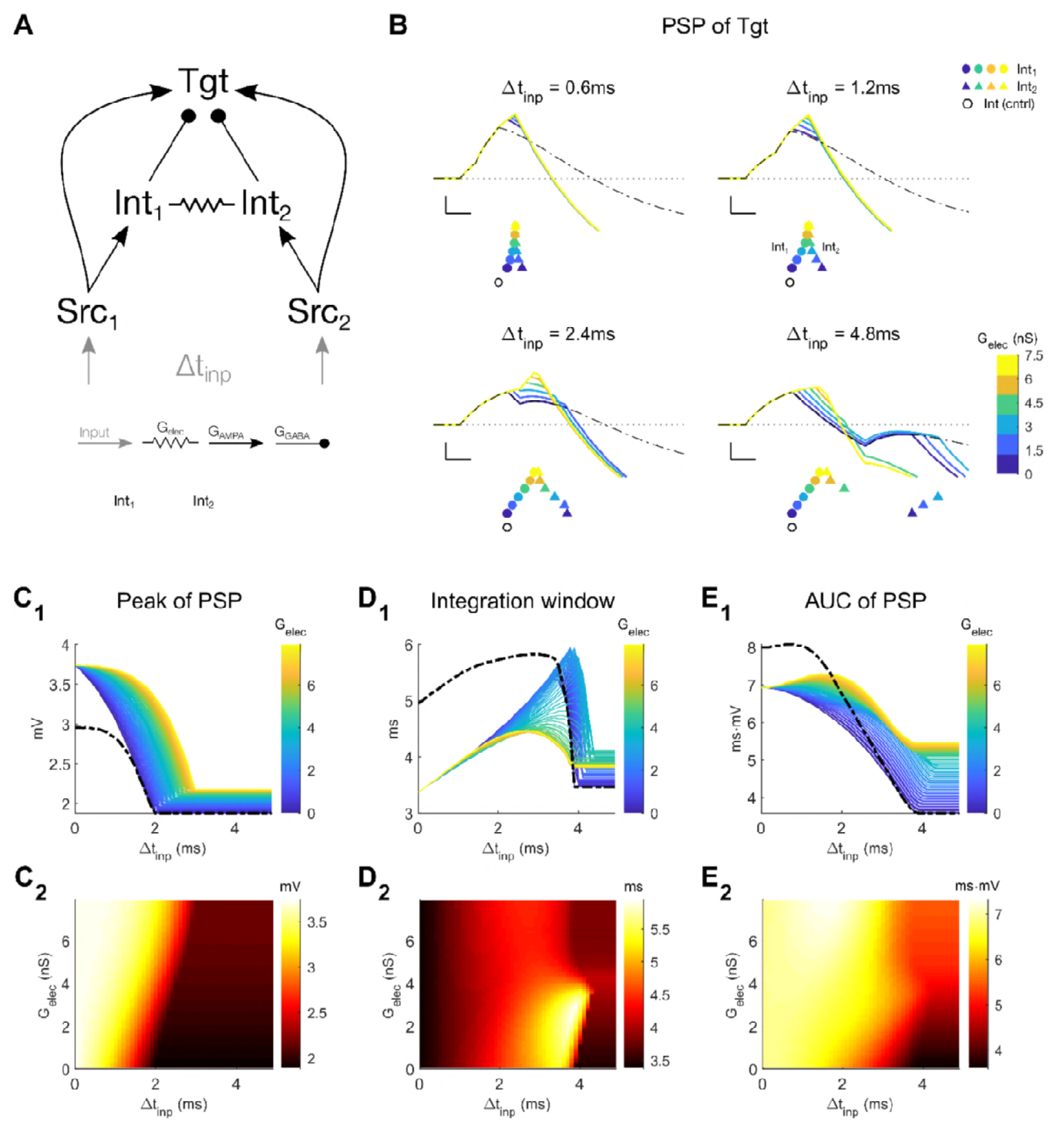
Model 2a: Two channels of Src, with two interneurons coupled by an electrical synapse. **A**: Model schematic; each Src neuron receives its own input, with timing difference between the two inputs as Δt_inp_. **B**: Examples of Tgt PSPs for different electrical synapse strengths between interneurons in the network (colored lines and legends). For comparison, the Tgt PSPs from Model 1b (Fig. 2), which has only one interneuron are also shown (black lines). Each subpanel represents different values of input timing differences Δt_inp_. Scale bar is 1 mV, 1 ms. Colored symbols represent the spike times of Int_1_ (circle •) and Int_2_ (triangle ▴); the black open circle (○) represents the spike time of the single Int in the control network (Model 1b). **C-E**: Increased electrical coupling leads to increased peak and AUC of the Tgt PSP, and decreased the integration window. In the top subpanels, colored lines represent different strengths of electrical coupling, while the black line represents the control network; the bottom subpanels show heat maps from the coupled network only.

We again provided this circuit with identical inputs, with time delay between the inputs Δt_inp_. With no electrical synapse, either channel of input from Src could separately drive a PSP in Tgt. In that case, similarly to Model 1b, summation of the two inputs in Tgt is limited to Δt_inp_ < 4 ms by the PSP (Fig. 3B, black lines). As electrical synapse strength increases, for Δt_inp_ < 4 ms we noted increased delays in Int_1_ spiking due to increased leak from the electrical synapse, but decreased delays in Int_2_ spiking due to the spikelet it received from Int_1_; together these effects result in a synchronizing effects of electrical coupling in this regime of input timing. Together, these changes in the timing of inhibition allowed for increased PSP peaks in Tgt (Fig. 3C), but dramatically decreased of the integration window in Tgt for Δt_inp_ between 2 and 4 ms (Fig. 3D), and increased the excited AUC (Fig. 3E). Hence, within Model 2a, electrical coupling enhanced Tgt input integration for specifically timed inputs, with increased PSP peaks within a narrowed integration window of the PSP. In the same circuit, for more than ∼4 ms of Δt_inp_, small increases in electrical synapse strength only served to increase leak in Int_2_ well after the spikelet had finished, ultimately delaying its spike. Larger increases in electrical synapse strength, however, allowed for the spikelet from Int_1_ to directly elicit spiking in Int_2_, which spiked earlier than it might have otherwise (Fig. 3B, lower right). The net effect allows the PSP in Tgt to increase by small amounts in peak (Fig. 3C) but the shortened integration window, resulting from the earlier spike in Int_2_, effectively prevents summation of the two Src inputs in Tgt. Thus, the varied effects of increased leak or excitatory spikelets resulting from an electrical synapse with varied strengths within even a simple circuit (Model 2a) increases flexibility for responses to signals passing through that circuit, compared to the same circuit with single interneuron (Model 1b).

A direct comparison between Model 1b, with one interneuron, and the most-strongly coupled results of Model 2a, is not apt, however: maintaining the synaptic parameters (see Methods) for Models 1b and 2a, meant that the total inhibitory conductance from Int to Tgt was doubled while the excitatory conductance from Src to Int was halved (see mismatch of dashed black lines and yellow lines in Fig. 3B). To explore a more consistent comparison of these two models, we ran the simulation again as Model 2a^norm^ with halved G_GABA→Tgt_ and doubled G_AMPA→Int_ (see Methods; Fig. 3S.A). Results from the strongly coupled 2a^norm^ were asymptotic to model 1b (yellow and dashed black lines Fig. 3S.B, C_1_ – E_1_). The trends from Model 2a^norm^ also preserved the trends from Model 2a for smaller values of Δt_inp_ and G_elec_ (Fig. 3S. C_2_ – E_2_). The larger differences in PSP peak and windows between Models 2a and 2a^norm^ occurred for weaker coupling and larger Δt_inp_, as the Tgt neuron in Model 2a^norm^ received smaller total inhibitory input compared to Model 2a (compare Fig. 3 with 3S). However, in this model the maximal G_elec_ in the range > 8 nS that would form the basis for a comparison with Model 1b exceeds the average physiological range of coupling coefficients (cc > 0.3).

While GABAergic coupling is rare between nearby electrically coupled inhibitory neurons of the thalamus [28, 32], it is sometimes observed between coupled pairs of inhibitory interneurons in cortex [29-31, 33]. To test the additional effects of GABAergic connectivity between electrically coupled interneurons, we included symmetrical GABAergic synapses in our model (Model 2b, Fig. 4A). From these simulations, we see that for transient inputs separated by Δt_inp_, the additional synapse further expanded the possibilities for subthreshold summation of inputs in Tgt. While the effect of G_GABA→Int_ on the peak PSP in Tgt was not substantially different from Model 2a, the integration window expanded with stronger inhibition (Fig. 4C), and the area under the excitatory part of the PSP was also expanded for stronger G_GABA→Int_ (Fig. 4D). Further, in the presence of stronger reciprocal inhibition, increased electrical coupling shifted the integration windows and AUCs to favor larger values of Δt_inp_ (Fig. 4C, D). Thus, electrical and inhibitory synapses compete for impact on the PSP.

**Figure 4.**
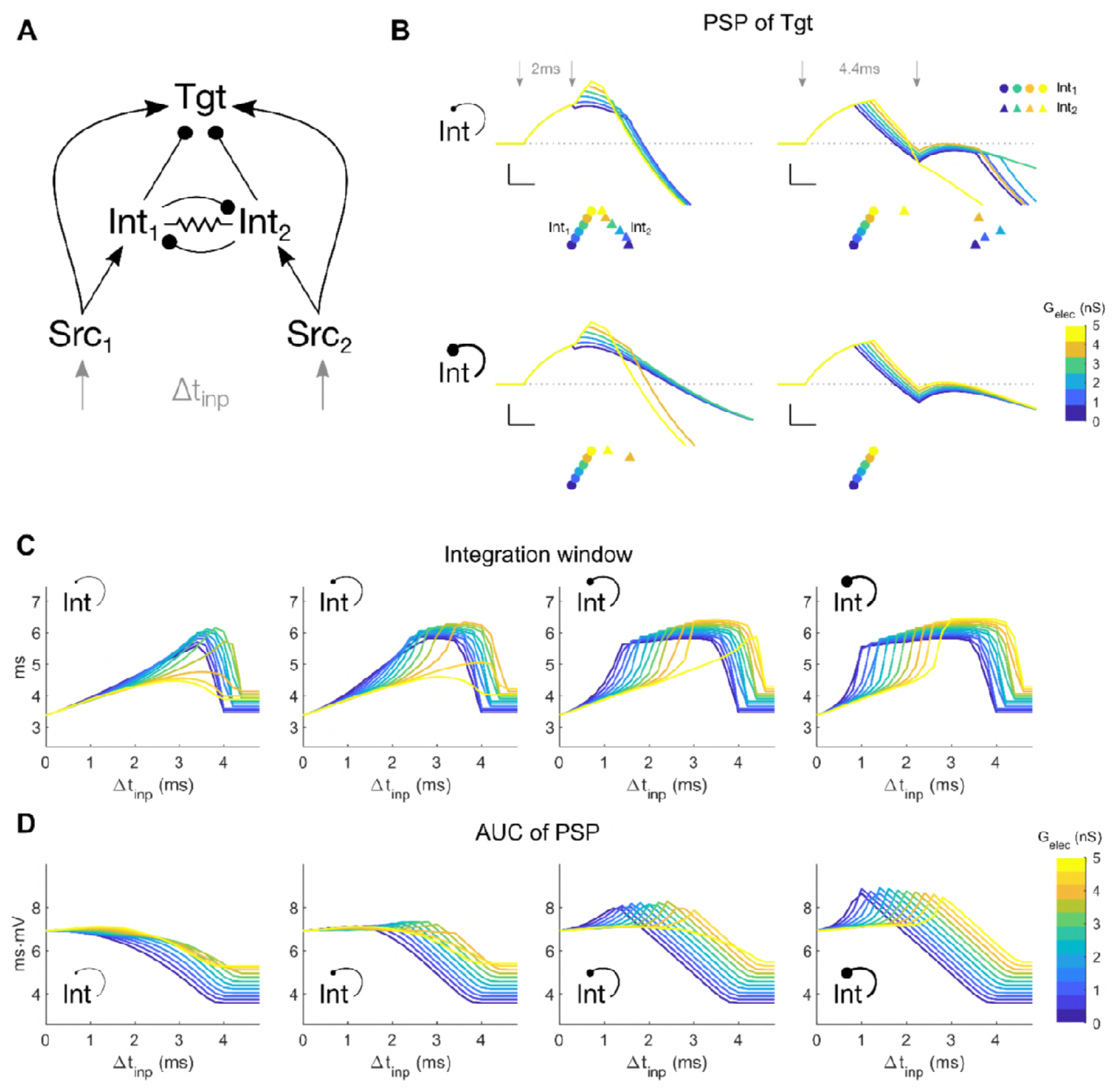
Model 2b: Two channels for Src, with both electrical and GABA coupling between the two Ints. **A**: Model schematic; each Src neuron receives its own input, with timing difference between the two inputs Δt_inp_. **B**: Examples of Tgt PSP for different electrical synapse strengths between interneurons of the coupled network (colored lines and legends). Each subpanel shows PSPs for different input timing differences Δt_inp_ (left: 2 ms, right: 4.4 ms), and strength of reciprocal inhibition 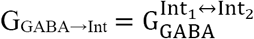 (top: 1 nS, bottom: 7nS). Scale bar is 1mV, 1ms. Colored symbols represent the spike times of Int_1_ (circles •) and Int_2_ (triangle ▴), with colors representing different values of G_elec_ between the two interneurons. **C-D**: Integration window and AUC of the PSP in Tgt for varied strengths of G_elec_ (0 – 5 nS) and G_GABA→Int_ (1, 3, 5, 7 nS from left to right).

Thus, similar to our previous demonstration [23], electrical synapse between inhibitory interneurons and their impact through inhibitory synapses onto a target work together in diverse ways to control the processing of transient signals passing through a circuit. We also note that changes in electrical synapse strength can potentially halve or double the PSP (Fig. 3C_1_), the integration window (Figs. 3D_1_, 4C) or area under the curve (Figs. 3E_1_, 4D); thus, modulation [5-7] or activity-dependent modifications of electrical synapses [9, 10] potentially have a powerful impact on summation of transient inputs in the Tgt cell.

### 2. Spiking responses in networks of canonical circuits with electrical synapses

To study a population of Tgt neurons of separate channels, we embedded 50 units of the canonical circuit into a network (Model Na, Fig. 5A), with electrical coupling between the Int neurons. To each Src neuron in the layer of 50, we provided identically sized inputs drawn from Gaussian distributions of input times with a standard deviation of σ_inp_ (Fig. 5B). In order to study spiking rather than subthreshold activity in the Tgt population, we increased G_AMPA_ from the Src to the Tgt (G_AMPA→Tgt_) and decreased G_AMPA_ from Src to Int (G_AMPA→Int_) in each unit, along with additional increases to Tgt excitability (see Methods) in order to elicit spiking in the Tgt neurons within 5-6 ms of Src spiking (consistent with latency to input in the somatosensory cortex [34]). We then quantified the distribution of spike times in the Int and Tgt populations (Fig. 5B). From these results, we observed that increases in electrical synapse strength acted to narrow the distributions of spike times in the Int layer (Fig. 5B, middle row), and markedly increased maximal amount of spiking for smaller σ_inp_. In the Tgt population, the narrowed Int distributions that resulted from increased electrical coupling acted to decrease the latency of Tgt from the input (Fig. 5B, bottom row and insets). Increased electrical synapse strength also decreased total spiking in Tgt, in fact selectively reducing later spikes, as a result of increased Int spiking.

**Figure 5.**
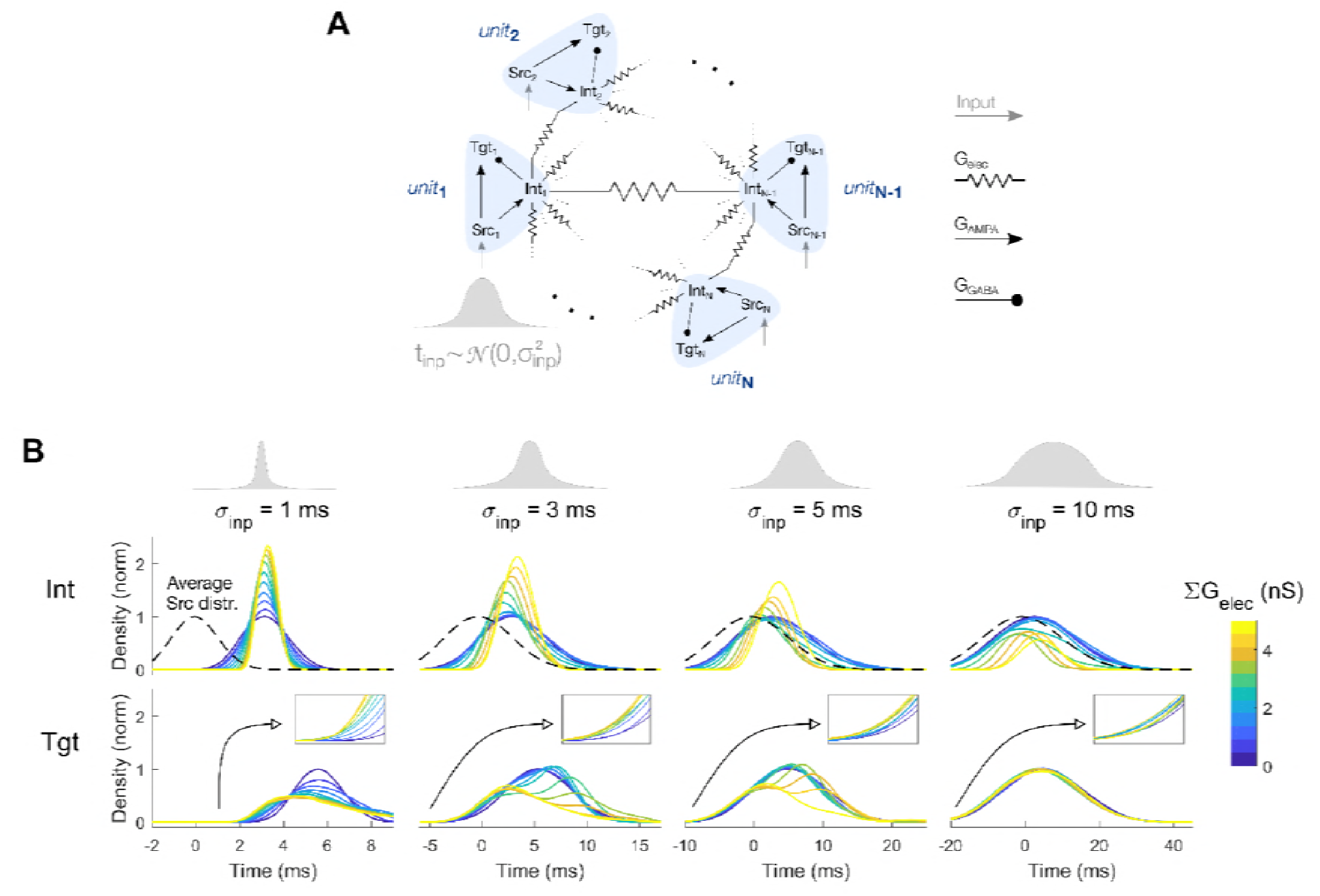
Model Na: Connected units of (Src, Int, Tgt) with interneurons connected by electrical synapses. **A**: Model schematic; each Src neuron receives a single input, with arrival times drawn from a Gaussian distribution with specific standard deviation σ_inp_. **B**: Normalized spike time distributions of the Src (top), Int (middle) and Tgt (bottom row) populations. Each column represents a different value of input timing distribution standard deviation (σ_inp_ = 1, 3, 5, 10 ms, from left to right). Dashed line (middle row) represents the average distribution of Src population spike times centered around t = 0ms. Insets (bottom row) show the latencies of Tgt population. Line colors represent different values of electrical coupling strength of the interneuron population.

Finally, we included GABAergic connectivity between the Ints of our network (Model Nb, Fig. 6A). The effects of electrical synapses on this network were similar to the previous model (Model Na): increases in electrical synapse strength decreased total spiking in Tgt, and selectively reduced later spikes, thus shifting its distribution to earlier times. Increased reciprocal inhibition was most effective for small values of σ_inp_, where stronger inhibition between Int neurons allowed Tgt neurons to spike more often, thus effectively counteracting the effect of electrical synapses (Fig. 6B, first two columns).

**Figure 6.**
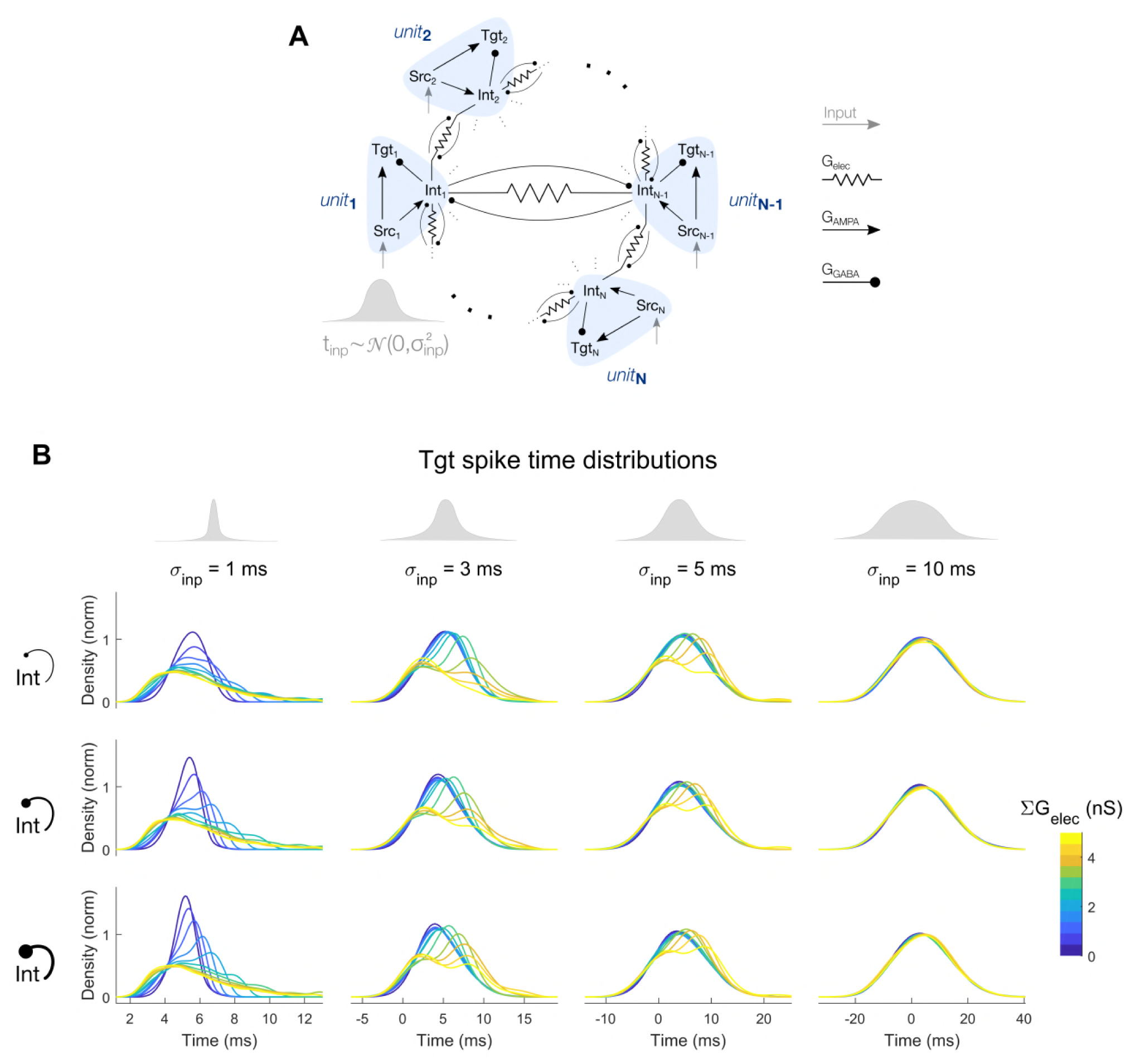
Model Nb: Connected units of (Src, Int, Tgt) with interneurons connected by electrical synapses and reciprocal inhibition. **A**: Model schematic; each Src neuron receives its own input, with arrival times drawn from a Gaussian distribution with specific standard deviation σ_inp_. **B**: Normalized spike time distributions of the Tgt population. Each subpanel represents a different combination of input timing distribution standard deviation (σ_inp_ = 1, 3, 5, 10 ms, from left to right) and reciprocal inhibition strength (ΣG_GABA→Int_ = 1, 3, 5 nS from top to bottom).

We compared our two network models by plotting the gain in spiking properties due to electrical synapses relative to the uncoupled case (ΣG_elec_ = 0) across input time distributions for the Int (Fig. 7A) and Tgt (Fig. 7B) populations. While the input was Gaussian, the Tgt distributions were often not Gaussian; therefore, we measured mean spike times, standard deviations of spike times, maximal density and total density of spiking, along with the relative latency. We observed that most of the effects that electrical synapses exerted on the output Tgt distribution were strongest for small σ_inp_. Mean spike times both increased and decreased for different combinations of σ_inp_ with inhibitory and electrical synapse strengths, while the spread (standard deviation) of spike times consistently increased with electrical synapse strength. Maximum density and total density of spiking, as well as relative latencies, decreased with increase in electrical synapse strength. Further, inclusion of larger reciprocal inhibition between the Int neurons led to decreased spiking within the Int population, thereby allowing Tgt neurons to spike faster, especially for the electrically uncoupled cases (Fig. 6B, dark blue lines). Increased electrical coupling combined with reciprocal inhibition led to increased inhibition within the Int layer, leading to more-synchronized Int activity but decreased total responses of the Int population. As seen previously, the effects of electrical and inhibitory synapses within the Int layer competed, such that Tgt spiking decreased less for stronger inhibition.

**Figure 7.**
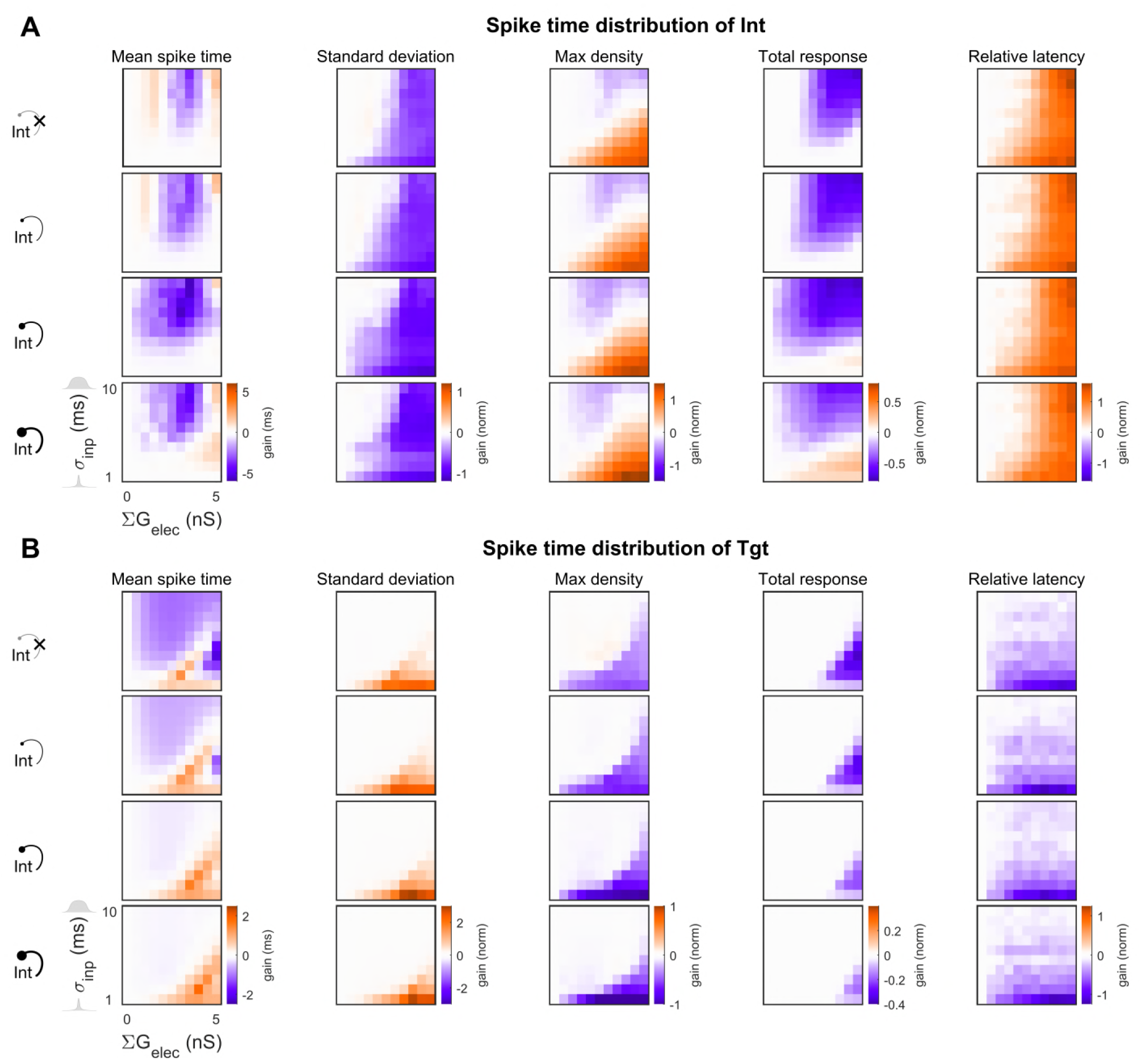
Properties of spike train distribution in Int and Tgt for Models Na and Nb. Values are expressed as normalized to the input (Src) distributions. (**A**) Int population and (**B**) Tgt population. In each panel, rows represent increasing reciprocal inhibitory strength within the Int population (ΣG_GABA→Int_ = 0, 1, 3, 5 nS, from top to bottom). The top row of each set is Model Na (Fig. 5), with ΣG_GABA→Int_ = 0 while the second, third and fourth rows represent Model Nb (Fig. 6), with ΣG_GABA→Int_ ≠ 0. The first column of each heat map always represents the uncoupled case, with 0 gain as indicated in white (see Methods). Within each heat map, electrical coupling ΣG_elec_ is varied on the x axis and input distribution size σ_inp_ is varied on the y axis.

Together, these results show that electrical synapses embedded within a network composed of canonical circuits have powerful and heterogeneous effects on the spiking of the Tgt output population, by altering spike times and total responses properties, as inputs from Src propagate through the network.

We quantified the mutual information between the spike time distributions of Src and Tgt, as well as the transmission efficiency from Src to Tgt (Fig. 8). For wider input distributions, we expected that mutual information would be larger due to increased entropy in both the input and output. Without any reciprocal inhibition within the Int layer, electrical coupling contributed to decreases in Src and Tgt mutual information, and more so with smaller input distributions (Fig. 8A, top row), as a result of dispersed Tgt spiking (cf. Fig. 6). Hence the transmission efficiency also decreased with electrical coupling, with more notable decreases with smaller σ_inp_ (Fig. 8B, top row). The largest decrease was roughly 35%.

**Figure 8.**
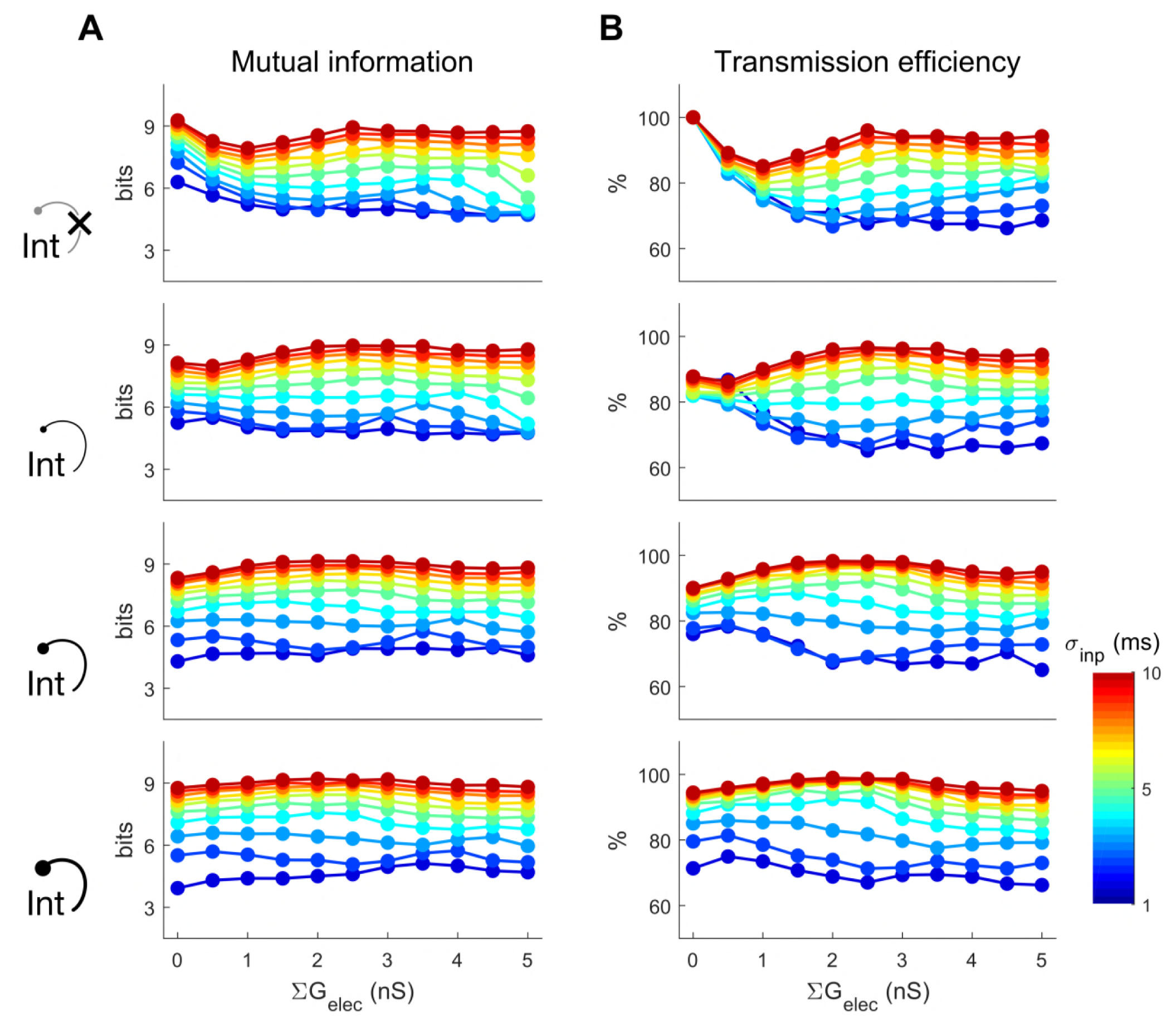
Information between the spike time distribution of Tgt and Src population in Model Na,b. **A**: Mutual information between Src and Tgt distribution; **B**: Transmission efficiency from Src to Tgt (% of Tgt entropy attributed to Src). Each row shows a different interneuron reciprocal inhibitory strength (ΣG_GABA→Int_ = 0, 1, 3, 5 nS from top to bottom). The line colors represent different input distribution standard deviations σ_inp_.

For the uncoupled case with no reciprocal inhibition, each Src elicited a single spike within its Tgt unit with predictable latency, leading to Tgt spike time distributions that mirrored Src distributions and resulted in maximal mutual information and 100% transmission efficiency (Fig. 8A&B, top row). As interneuron reciprocal inhibition was added and spiking in the Int population decreased, some neurons within the Tgt distribution were able to spike much faster but with less uncertainty, creating a smaller distribution (Fig. 6B) and smaller entropy; as a result, reciprocal inhibition led to decreases in both mutual information and transmission efficiency.

## Discussion

Overall, our simulations show that electrical synapses between interneurons in canonical networks regulate both subthreshold activity and network spiking activity, ultimately exerting powerful effects on the output activity of the network while processing and passing on its inputs. The general effect was, for closely-timed inputs, that increase in electrical coupling strength led to (a) the delay of spiking in the inhibitory interneurons, which enabled larger summations in the target output earlier; yet simultaneously (b) stronger synchronized inhibition from the interneuron population to the target population at a later time, which limited the output of target. These competing effects highlight the diverse roles that electrical synapses of dynamically varying strength might play in circuits across the brain.

Electrical synapses in a local network regulate subthreshold summation of inputs in the target neuron. Stronger electrical coupling allowed the target neuron to integrate its source inputs with higher summed PSP peaks, yet limited time windows for further inputs to summate. Furthermore, changes in electrical coupling in a local network of interneurons, possibly via electrical synapse plasticity, led to more flexibility in regulating subthreshold summation than a global inhibitory neuron with varied excitability. However, our results also showed that reciprocal inhibition between the electrically coupled interneuron pair expanded the integration window and the area under the curve of the target PSP, especially for relatively larger differences in input timings. This suggests that the competition between the electrical coupling and reciprocal inhibition within the local interneuron networks could regulate the ability for the target neuron to either be a coincidence detector or an integrator.

At a network level, we find that electrical coupling of the interneuron population modulates the target population activity over different distribution of input timings. Similar to the subthreshold effect, increase in electrical synapse strength led to a more delayed, yet more concentrated activity in the interneuron population, effectively synchronizing their activity. Hence, stronger electrical coupling allowed stronger earlier response of the target layer activity but weaker response afterwards. However, because the activity of the interneurons was limited within a smaller temporal window due to electrical coupling, inhibition towards the target population was limited in time, hence the output activity was more sustained compared to uncoupled cases. As a result, electrical coupling allowed earlier yet more spread out responses and effectively reduced both the mutual information and the transmission efficiency, but mainly for small input distribution sizes. One implication of this is, although corrupting the integrity of input-output temporal coding, electrical coupling between the interneurons could increase temporal heterogeneity as inputs coming from different sources arrive too closely with each other. On the contrary, reciprocal inhibition within this population decreased the interneuron activity, which led to much less decrease of target response in presence of electrical coupling. However, for closely-timed input distributions, the target temporal code distribution was also changed towards less output timing spread, especially for electrically uncoupled or weak coupled cases, resulting in loss of mutual information and transmission efficiency in the presence of reciprocal inhibition.

These results point toward yet another competition between electrical coupling and reciprocal inhibition within the interneuron population at the network level to regulate the temporal code of the output distribution. Although both disrupt the input-output temporal integrity of closely-timed input distributions, electrical synapses increase temporal heterogeneity while the inhibitory synapses increase temporal homogeneity. As transient processing due to electrical coupling of inhibitory neurons does not receive much attention, the implications are not known. As recent work shows, there exist gap junctions between PV interneurons across barrel boundaries [35], suggesting there may be electrical coupling even across different barrel cortex via intermediate interneurons. One implication of our network simulations is that temporal heterogeneity due to electrical coupling would help separate inputs coming too close to each other in time but from different sources.

## Methods

### 1. Model and simulation

The canonical disynaptic feedforward-inhibition network can be simplified as a small 3-cell network (Fig. 1A_1_), comprising of a Src (source), an Int (interneuron) and a Tgt (target) neuron. For subthreshold investigations, we explored small networks with either 1 Int (Models 1a and 1b), 2 Ints (Models 2a, 2a_norm_ and 2b). For activity explorations, we used Models Na and Nb (containing N Ints neurons).

Coincidentally, N also equals the number of units, each of which is Model 1a. In this paper, we used N = 50.

#### 1.1 Izhikevich type neuron model

For generalizability, we modelled Src and Tgt as regular spiking (RS) neurons and Int as a fast spiking (FS) neuron with Izhikevich formulism [36]. Briefly, eqs. 1-2 describe the dynamics of the membrane potential *v* and recovery current *u* respectively, with the spiking condition in eq. 3. Additionally, implementation of FS neuron model also differs from RS as described in eqs. 4-5 [36].

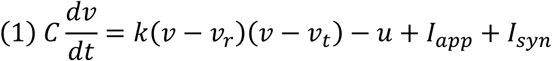

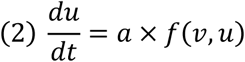

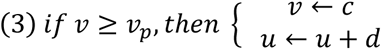

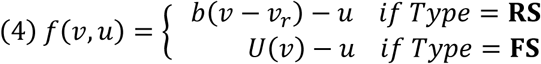

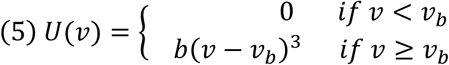

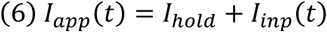

We applied a holding current I_hold_ (eq. 6) of 50 pA to Int to easily evoke spiking in response to input from Src, except for the variations of I_hold_ to Int from an array of [0, 25, 50, 60] pA to account for different Int excitability in Model 1b, Fig. 2. For subthreshold investigation (Models 1a-2b), we modelled Src and Tgt with the same set of parameters of an RS neuron (Table 1). To inspect network activity (Models Na, Nb), we tuned the parameters (halved capacitance and lowered threshold potential) and applied a 10 pA holding current for each Tgt neuron in order for its Src to easily evoke its spiking.

**Table 1.** Izhikevich model parameters. Single asterisk * is for Models 1a-2b. Double-asterisk ** is for Models Na, Nb.

|  | Src | Int | Tgt |
| --- | --- | --- | --- |
| Type | RS | FS | RS |
| $C$ (pF) | 100 | 20 | 100* or 50** |
| $v_r$ (mV) | -60 | -55 | -60 |
| $v_t$ (mV) | -40 | -40 | -40* or -45** |
| $v_p$ (mV) | 35 | 25 | 35 |
| $k$ (nS) | 0.7 | 1 | 0.7 |
| $a$ (1/ms) | 0.03 | 0.2 | 0.03 |
| $b$ (nS) | -2 | 0.025 | -2 |
| $c$ (mV) | -50 | -45 | -50 |
| $d$ (pA) | 100 | 0 | 100 |
| $v_b$ (mV) | | -55 | |

#### 1.2 External input to Src neurons

In all cases, only Src received external input: a brief 20-30 ms of 200-300 pA DC input, sufficiently to evoke a single action potential in in Src (eq. 7). For Models 1b, and 2a, a^norm^, b, we varied the arrival time differences between input to Src_2_ and input to Src_1_ as Δt_inp_ from 0 to 20 ms. For Models Na and Nb, timings of Src inputs were drawn from a normal distribution with standard deviation as σ_inp_ in which we varied from 1 to 10 ms.

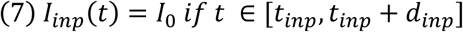

#### 1.3 Synaptic connections and different network configurations

For synaptic inputs, neurons can either excite each other via AMPA synapses, inhibit each other via GABA synapses or couple with each other via electrical synapses, as described in eqs 8-11. Src sends AMPA excitatory input to Tgt and Int separately sufficiently to drive Int to spike and for Tgt to receive a noticeable EPSP (Models 1a – 2b) or to spike (Models Na, Nb). Int sends GABAergic inhibitory input to Tgt. In our simulations, Ints are either only electrically coupled (Models 2a, 2a^norm^ and Na) or both electrically coupled and reciprocally inhibit each other (Models 2b, Nb).

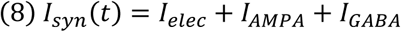

Electrical synapses were implemented as symmetric linear resistance, as shown in eq. 9. For two coupled Int neurons, we varied the electrical synapse conductance of from 0 - 8 nS (unless otherwise noted), corresponding to coupling coefficients (cc.) of roughly 0 – 0.33. For larger coupled networks (Models Na,b), Ints are electrically coupled homogeneously in an all-to-all manner (Fig. 5A, 6A), with each coupling conductance scaled to the number of Ints as G_elec_ = ΣG_elec_ /N_Int_.

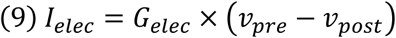

Chemical synapses were implemented with a single exponential decay as described in eqs. 10-11, and implemented following the example of [37] in *Brian2* documentation. The synaptic reversal potentials and time constants were fixed: E_AMPA_ = 0 mV, τ_AMPA_ = 2 ms & E_GABA_ = −80 mV, τ_GABA_ = 10 ms. The conductance parameters were either fixed or varied as in Table 2. The difference between model 2a and 2a^norm^ was the synaptic conductances to Int and Tgt: the former has similar parameters as model 1b, whereas the latter has G_AMPA→Int_ doubled and G_GABA→Tgt_ halved to account for the existence of the additional Int_2_ neuron, in order to compare between Model 1b and the highly coupled configuration of Model 2a. For model Nb where, in addition to electrical coupling, the Int population also reciprocally inhibits itself in an all-to-all manner; each inhibitory conductance was also scaled to the number of Int’s as G_GABA→Int_ = ΣG_GABA→Int_ /N_Int_.

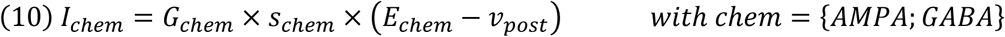

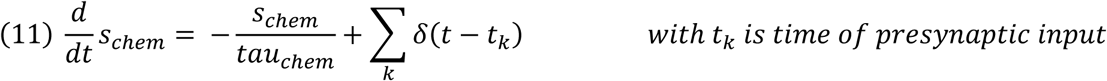

**Table 2.**
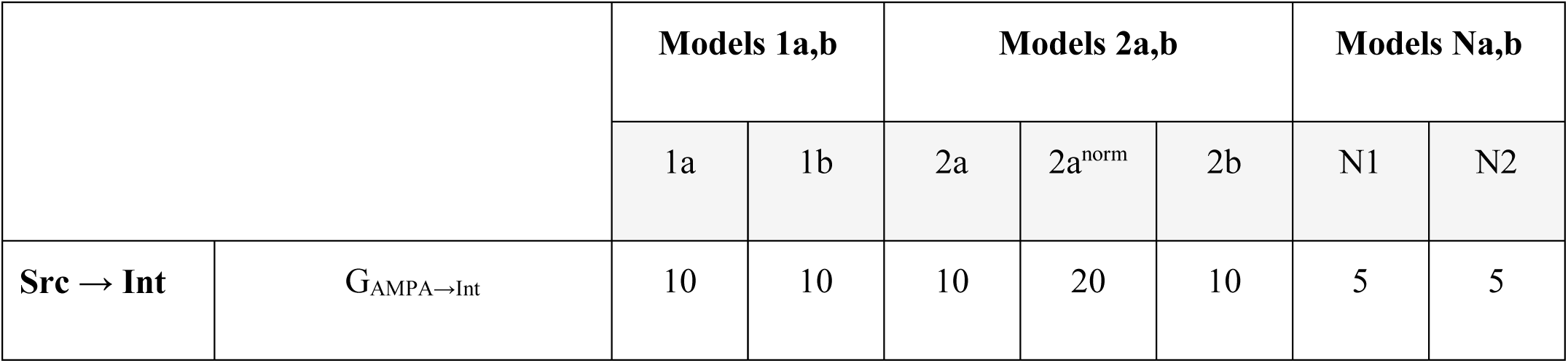
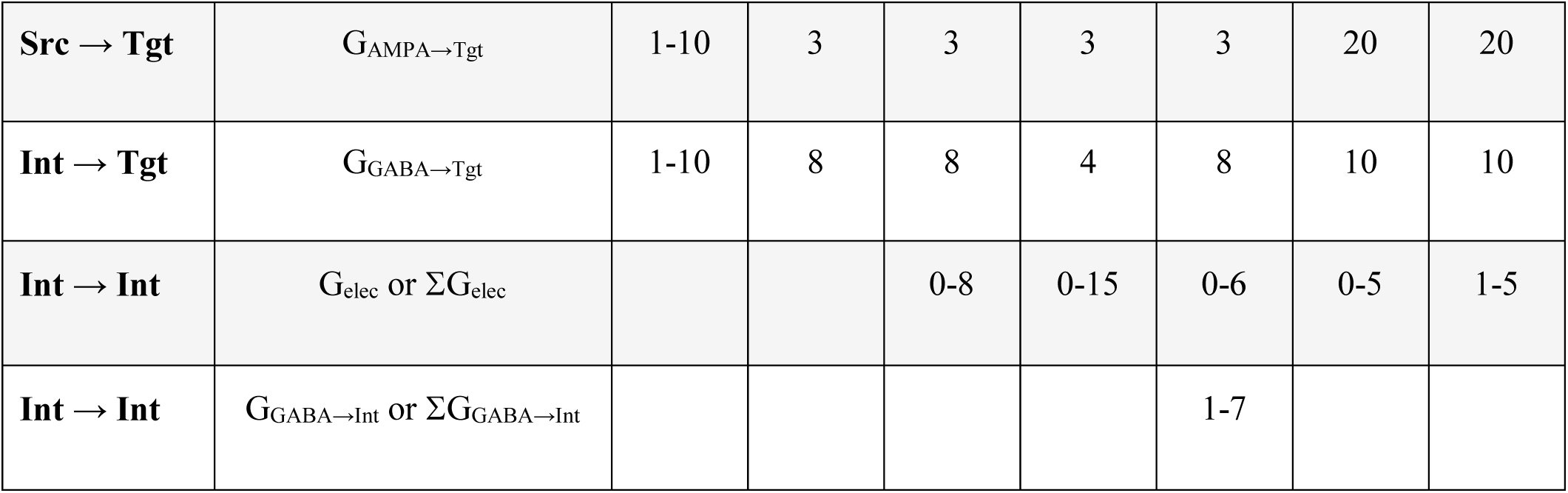
Synaptic conductances for different network configurations.

|  |  | Models 1a,b |  | Models 2a,b |  |  | Models Na,b |  |
| --- | --- | --- | --- | --- | --- | --- | --- | --- |
|  |  | 1a | 1b | 2a | 2a<sup>norm</sup> | 2b | N1 | N2 |
| Src → Int | $G_{AMPA \rightarrow Int}$ | 10 | 10 | 10 | 20 | 10 | 5 | 5 |
| **Src → Tgt** | $G_{\text{AMPA} \rightarrow \text{Tgt}}$ | 1-10 | 3 | 3 | 3 | 3 | 20 | 20 |
| **Int → Tgt** | $G_{\text{GABA} \rightarrow \text{Tgt}}$ | 1-10 | 8 | 8 | 4 | 8 | 10 | 10 |
| **Int → Int** | $G_{\text{elec}}$ or $\Sigma G_{\text{elec}}$ | | | 0-8 | 0-15 | 0-6 | 0-5 | 1-5 |
| **Int → Int** | $G_{\text{GABA} \rightarrow \text{Int}}$ or $\Sigma G_{\text{GABA} \rightarrow \text{Int}}$ | | | | | 1-7 | | |

#### 1.4 Simulation environment

Simulations were run in the Python-based open source simulator *Brian2* [38]. Subthreshold simulations were run for 100 ms with dt = 0.01 ms (Models1a-2b). For each parameter set in network activity investigation (Models Na, Nb), 50 random simulations were run as external input timings to Src population were drawn from a normal distribution with size σ_inp_, in which each simulation was 200 ms and dt = 0.05 ms because less accuracy was required and for speed.

### 2. Analysis

Analysis and visualization were mainly performed in MATLAB (MathWorks R2018a) and the open source graphics editor Inkscape 0.92.3.

#### 2.1 Subthreshold investigation

For subthreshold investigations (Fig. 1 – 4), we obtained the net postsynaptic potential (PSP) of the Tgt neuron and quantified the peak potential, duration (or integration window) and area under the curve (AUC) of the positive portion of the PSP (Fig. 1B).

#### 2.2 Network activity investigation

For each set of parameter θ, we obtained the raw distribution of spike times X(θ, *C*) = {X_*k*_(θ, *c*_*i*_)} population *C* aggregated from all X_*k*_(θ, *c*_*i*_), which is the spike time array of neuron *c*_*i*_ in simulation *k*^*th*^. The symbol *C* (or *c*) represents the population name, can either be any of the following {Src, Int, Tgt}. *i* = {1, 2 … *N*_*C*_} with *N*_*C*_ as the number of neurons in population *C*. *k =* {1, 2 … *N*_*s*_} with *N*_*s*_ as the number of random simulations. In this paper, we used *N*_*C*_ = 50 with all *C* and *N*_*s*_ = 50 as described earlier.

##### 2.2.1 Normalized properties of spike time distributions

To easily compare between different initial input distributions, we generally normalized all quantifications to the Src population (Fig. 5 – 7). More specifically, for each X_*C*_ = X(θ, *C*), we defined normalized mean spike time as the difference between the mean of X_*C*_ and that of X_*Src*_. The normalized standard deviation was the standard deviation of X_*C*_ normalized over the standard deviation of X_*Src*_.

For each X_*C*_ = X(θ, *C*), we calculated the spike density from the smoothed histograms of spikes times. More specifically, each array of spike times X_*C*_ was histogrammed with a bin width that equals to one-tenth of the σ_inp_ in order to avoid under-sampling with small σ_inp_ and over-sampling with large σ_inp_; then it was smoothed by convoluting with a Hanning window of size 20 to obtain the un-normalized density *d*_*C*_(t). For visualization, the spike times were translated relative to the mean Src spike time distributions, whereas the densities were scaled over the maximum density of the Src distribution to calculate the normalized density *D*_*C*_(t). Note: neither *D*_*C*_(t) nor *d*_*C*_(t) represented estimated probability density function, because the smoothed histograms were not normalized by their number of samples.

For quantification comparison, we defined normalized maximum density as the maximum density of *d*_*C*_(t) normalized over that of *d*_*Src*_(t). The normalized total response was calculated by normalizing the area under the curve of *d*_*C*_(t) over that of *d*_*Src*_(t) (note: neither *D*_*C*_(t) nor *d*_*C*_(t) represented estimated probability density function, hence AUC was not necessarily 1). Lastly, the relative latency was defined as the time point which *d*_*C*_(t) reached 10% of maximum Src density, scaled by the standard deviation of Src spike time distribution X_*Src*_.

Additionally, gain of a particular property *Q* of a spike time distribution due to a parameter set θ was defined as the difference between itself and the same property when the electrical coupling parameter in set θ equals to 0, in other words *Gain*[*Q*(θ)] = *Q*(θ) – *Q*(θ_electrically uncoupled_).

##### 2.2.2 Mutual information and transmission efficiency

For network investigation, we also quantified the mutual information and transmission efficiency between the Src and Tgt population spike time distribution (Fig 8). Here we considered Src to be an input channel, whereas Tgt to be an output channel.

For each X_*C*_ = X(θ, *C*), we estimated the probability function *p*(*C*) by histogramming the spike time arrays X_*C*_ with a fixed bin width of 0.01ms. The joint probability function *p*(*Src, Tgt*) of Src and Tgt was also estimated by histogramming all the spike time pairs of (X_*Src*_ X_*Tgt*_) with similar bin widths. We consider any missing spike (for example with cases that Src_i_ did not induce any spike in Tgt_i_ due to certain network configurations or parameter set) to take the value of max(X_*C*_) + 2σ(X_*C*_) to account for more accurate estimation of the marginal distribution of both Src and Tgt. Taking these cases out led to misrepresentation of the marginal distribution and join distribution.

We calculated the mutual information between Src and Tgt with eq. 12 in which H(*A*) is the entropy of the distribution *p*(*A*) (eq. 13) and H(*A, B*) is the entropy of the joint distribution *p*(*A, B*) (eq. 14).

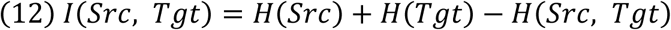

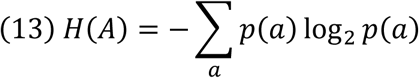

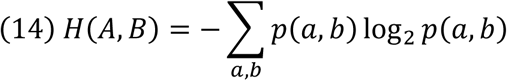

We measured the transmission efficiency from the input channel (Src) to the output channel (Tgt) with eq. 15 [39]. This could be interpreted as % of the entropy of output that could be attributed to the input.

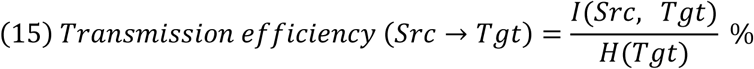

All code is available upon reasonable request.

## Acknowledgements

Parv Venkitasubramaniam for helpful discussions; Funding from NSF IOS 1557474, Whitehall Foundation

---

**Figure 3 Supp.**
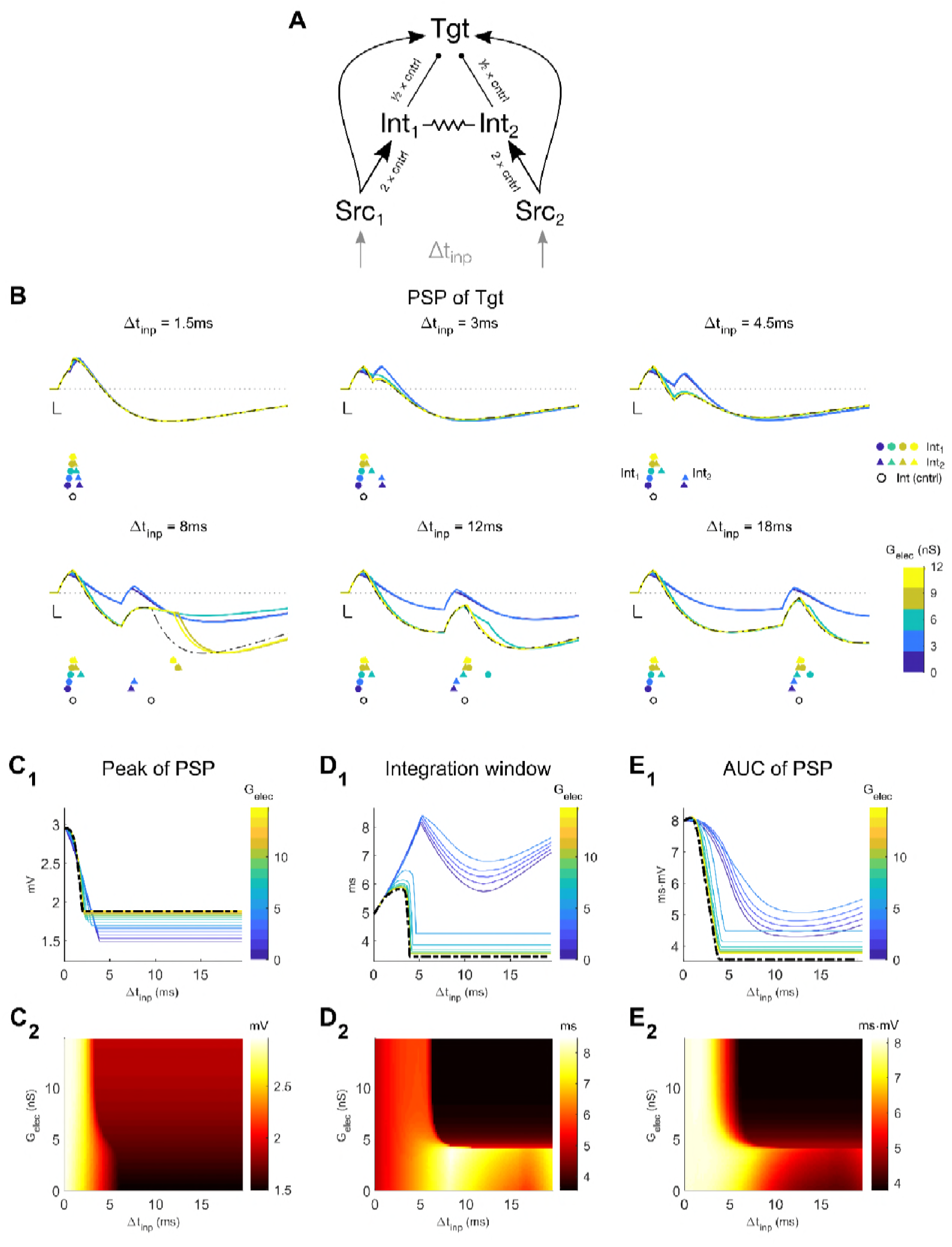
Model 2a^norm^: Comparison between single-interneuron control network and coupled interneuron network with normalized synaptic strengths. **A**: Schematic for the network identical to Model 2a but with 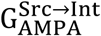 doubled and 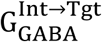 halved. This was done to easily compare between the coupled network and the control network. **B**: Examples of Tgt PSP due to different electrical synapse strengths between interneurons of the coupled network (colored lines and legends), in comparison to the control single-interneuron network (black dashed-dotted lines and legends). Each subpanel represents different values of input timing differences. Scale bar is 1 mV, 1 ms. Colored legends represent the spike times of Int_1_ (filled circle •) and Int_2_ (filled triangle ▲), in which each color represents different values of G_elec_ between the two interneurons in the coupled network. The black open circle (○) represents the spike times of the single Int neuron in the control network. The dotted line represents Tgt resting potential. **C-E**: The effects on integration window of the coupled network asymptote those in the control network with sufficiently large interneuron electrical coupling, although corresponding to highly unlikely coupling coefficient. For each panel, subpanel *1* (top) shows 2D plot with each colored line representing different electrical coupling, while the dashed-dotted line represents the control network; subpanel *2* (bottom) shows the heatmap of the same data from the coupled network.

